# Juvenile hormone, but not nutrition or social cues, affects reproductive maturation in solitary alkali bees (*Nomia melanderi*)

**DOI:** 10.1101/134387

**Authors:** Karen M. Kapheim, Makenna M. Johnson

**Author notes:** Correspondence: Karen M. Kapheim.

## Abstract

Eusocial insect colonies are defined by extreme variation in reproductive activity among castes, but the ancestral conditions from which this variation arose are unknown. Investigating the factors that contribute to variation in reproductive physiology among solitary insects that are closely related to social species can help to fill this gap. We experimentally tested the role of nutrition, juvenile hormone, and social cues on reproductive maturation in solitary alkali bees (Halictidae: *Nomia melanderi*). We find that alkali bee females emerge from overwintering with small Dufour's glands and small ovaries, containing oocytes in the early stages of development. Oocyte maturation occurs rapidly, and is staggered between the two ovaries. Lab-reared females reached reproductive maturity without access to mates or nesting opportunities, and many had resorbed oocytes. Initial activation of these reproductive structures does not depend on pollen consumption, though dietary protein or lipids may be necessary for long-term reproductive activity. JH is likely to be a limiting factor in alkali bee reproductive activation, as females treated with JH were more likely to develop mature oocytes and Dufour's glands. Unlike for related social bees, the effects of JH were not suppressed by the presence of older, reproductive females. These results provide important insight into the factors that influence reproductive activity in an important native pollinator, and those that may have been particularly important in the evolution of reproductive castes.

## Introduction

Complex social organization, such as that observed among honey bees (*Apis mellifera*), ants (Formicidae), and vespid wasps (Vespidae), is marked by a high degree of variance in reproductive activity among individuals within a colony. This variation is demarcated among reproductive castes, whereby workers do not reproduce, despite engaging in maternal behaviors (e.g., brood feeding or nest defense), and queens reproduce while largely refraining from brood care (Michener, 1974). Workers of many social insect species have similar reproductive anatomy to queens (e.g., ovaries, spermathecae, glands, ovipositor), yet remain functionally sterile. This suggests the factors that influence the function of these structures differ between queens and workers, and that understanding this variation may provide insights into the physiological basis for the origin of social insect castes.

The factors that differentially influence reproductive activation among social insect castes include nutritional, endocrine, and social cues (Kapheim, 2017). For example, both queens and workers acting as nurses within a honey bee colony consume a protein-rich diet, but this protein contributes to egg-production only in the queens (Winston, 1987). Similarly, treatment with the juvenile hormone (JH) analog methoprene leads to accelerated ovarian development in queen paper wasps (*Polistes canadensis*), but instead increases foraging activity in workers (Giray et al., 2005). Finally, social cues, such as aggression from the queen, can repress endocrine pathways, and thus ovary maturation, in worker bumble bees (*Bombus impatiens*) and social halictid bees (*Megalopta genalis*), but aggression directed from workers toward queens does not have the same effect (Kapheim et al., 2016; Padilla et al., 2016; Smith et al., 2009). Understanding how these differences in sensitivity to physiological and environmental cues arise among females and contribute to variation in reproductive activity is thus key to understanding the origins of social insect castes.

One approach toward this goal is to investigate how these factors contribute to variation in reproductive development in solitary species representative of the ancestors that gave rise to social castes. We conducted two experiments to determine how variation in nutrition, JH, and the social environment influence reproductive development in a solitary bee that shares similarities with the ancestors of social bees. Alkali bees (*Nomia melanderi*) are semi-managed, native pollinators of alfalfa seed crops that range throughout the western U.S.A. (Cane, 2008). This species belongs to a basal subfamily of Halictidae (Nomiinae) in which eusociality has never evolved, but they are closely related to the Halictinae, in which social behavior is highly variable (Michener, 1974; Michener, 2007), and eusociality evolved at least twice (Brady et al., 2006; Danforth et al., 2008; Gibbs et al., 2012). Bees in the family Halictidae shared a common ancestor with the Apidae (e.g., bumble bees, honey bees) approximately 115 mya, and may thus have important differences in reproductive physiology (Cardinal and Danforth, 2013). Alkali bees are considered solitary, because each female has her own nest, but they often nest in close proximity to other females in large aggregations (Cane, 2008; Wcislo and Engel, 1996). Importantly, these bees exhibit extended maternal behavior, characteristic of that which was a necessary pre-adaptation to sociality (Batra, 1970; Batra and Bohart, 1969; Schwarz et al., 2007). As such, understanding the factors that influence reproductive physiology in this species can shed light on the physiological basis for social evolution.

Little is known about the factors that influence alkali bee reproductive development. However, a recent study demonstrated that JH accelerates reproductive maturation (Kapheim and Johnson, 2017). It was also recently documented that adult female alkali bees consume pollen on a daily basis (Cane et al., 2016). Pollen is the primary source of dietary protein and lipids for bees (Roulston and Cane, 2000), but whether pollen consumption is necessary for reproduction has not been experimentally tested. We investigated two aspects of reproductive physiology – oocyte growth, which requires proteins for egg-yolk (Badisco et al., 2013) and maturation of the Dufour's gland, which secretes lipids used for nest cell construction (Cane, 1981). Our results reveal that vitellogenesis can occur rapidly among newly emerged females, even without mating or nesting opportunity. The initiation of oogenesis and Dufour's gland maturation does not require dietary protein, and females treated with JH were more likely to reach reproductive maturity. This response to JH was not affected by variation in the social environment (i.e., co-housing with an older, reproductive female). This provides important insight into the physiological foundation from which social insect castes evolved, as well as the reproductive physiology of an important pollinator.

## Methods

### Collections

This study took place in Touchet, WA, U.S.A, where alkali bees nest in large soil beds near alfalfa seed fields (Cane, 2008). Alkali bees overwinter as prepupae in below-ground nests, and emerge as adults upon completion of development the following summer. We trapped newly eclosed adult females from 27 May-8 June 2016 by placing emergence traps on 3 bee beds prior to the start of emergence. Traps were checked at least 3 times a day, and new bees were transferred back to the laboratory in individually labeled 15 ml tubes placed inside a cooler with a single ice pack placed under a layer of cardboard.

### Experiment 1 – nutrition effects on reproductive physiology

Upon arrival to the laboratory, bees were chilled at 4ºC for 5 min and randomly assigned to a treatment group: sugar water only (sterile 35% sucrose solution), sugar water with pollen (2.5 g sterile, finely-ground, honey-bee pollen in 30 ml of sterile 35% sucrose solution), sugar water with pollen plus 4 sprigs of fresh, untripped alfalfa flowers. (Bees were observed manipulating these flowers to release pollen on a regular basis throughout the experiment.) Bees were placed in perforated plastic deli containers (72 mm x 90 mm lower diameter x 113 mm upper diameter), and reared in the lab for 10 days (d). Sugar water or pollen mix and alfalfa flowers were changed daily. Pollen-sugar mixture was shaken vigorously before each feeding to achieve homogeneity, and then pipetted into feeding troughs made from 1.5 ml microcentrifuge tubes with the tapered tip removed. The cages were kept at 22-28ºC, 40-85% RH and full spectrum lighting 13 L: 11 D, as has been previously described (Kapheim and Johnson, 2017). At the end of the 10 d rearing period, bees were chilled for 3 min at 4ºC, placed in individually-labeled tubes, and flash-frozen in liquid nitrogen.

We also collected newly emerged females and reproductive females of unknown age for comparison to lab-reared females. Newly emerged females were collected from emergence traps as described above, but were flash-frozen immediately upon return to the laboratory. Reproductive females were identified as those returning to a nest hole with pollen on their hind legs. They were captured by net, and flash-frozen immediately upon return to the laboratory.

### Experiment 2 – social and endocrine effects on reproductive physiology

Newly emerged females were collected as in Experiment 1. Each bee was randomly assigned to a treatment group: sham control, solvent control, or JH. For JH treatments, JH-III (product E589400, Toronto Research Chemicals, Inc., Toronto, Ontario, Canada) was dissolved in dimethylformamide (DMF) at a concentration of 50 μg per μl. Bees in the solvent control group received 1 μl of DMF applied to the thorax with a pipette tip. Bees in the JH group received 50 μg JH in 1 μl of DMF applied to the thorax with a pipette tip. Bees in the sham group were touched lightly on the thorax with a clean pipette tip. Hormone treatments were repeated when bees were 5 d old. Bees in each treatment group were randomly assigned to be caged alone or with an older, reproductive female, defined as above. All bees were paint-marked on the dorsal abdomen with a uniquely colored Decocolor^©^ paint pen (Uchida of America Co, Torrance, CA). All bees were reared in cages, and received 35% sugar water mixed with pollen and fresh alfalfa flowers for 10 d, as described for Experiment 1.

Upon collection, all bees from both experiments were stored in liquid nitrogen until return to Utah State University, where they were transferred to a −80 ºC freezer.

### Dissections

Dissections followed previously reported methods (Kapheim and Johnson, 2017). Briefly, bees were dissected under a Leica M80 stereomicroscope fitted with an IC80HD camera (Leica Microsystems, Buffalo Grove, IL, USA). We measured Dufour's gland and terminal oocyte lengths from images, using software in the Leica Application Suite (v. 4.5). The observer was blind to the treatment group of each bee during dissections, and both authors concurred on measurements. Mating status was determined by examination of the spermatheca under a compound microscope. We excluded newly emerged females with sperm and reproductive females without sperm from further analyses.

### Stages of ovary maturation

Images were further analyzed to classify ovaries into stages of oocyte maturation. Like most halictid bees, alkali bees have three ovarioles in each of two ovaries. The following categories were modified from Duchateau and Velthuis (1989) and Oliveira et al. (2017). Stage I – the oocyte and associated trophocytes occupying a single egg chamber can be distinguished from one another, but the oocyte is much smaller than the trophocytes and is spherical in shape. Stage II – the oocyte is smaller than the trophocytes, and is cylindrical in shape rather than round. Stage III – the trophocytes and oocyte are similar in size, the oocyte occupying 45-55% of the length of the egg chamber. The oocyte is elongated. The trophocytes appears less opaque, and the oocyte appears more solid and full. It is during this stage in which vitellogenesis is initiated. Stage IV – the oocyte is much longer than the trophocytes, and occupies more than 55% of the total egg chamber. The trophocytes appear translucent and small in the anterior portion of the egg chamber. Stage IVr (reabsorbing) – the oocyte has all the characteristics of stage IV, but is misshapen and yellow in color, indicating reabsorption. Stage V – the oocyte is large, robust, and opaque with no associated trophocytes. Vitellogenesis is complete at this stage. Stage Vr (reabsorbing) – oocyte has no trophocytes present, but has the characteristics of being reabsorbed. When possible, we measured the length of the maximum terminal oocyte and its associated trophocytes of both resorbing (if present) and viable oocytes, and used these measurements to classify stage of ovary maturation. We identified the maximum terminal oocyte and maximum stage of oocyte maturation as the longer/more mature from the two ovaries. We used these maxima for statistical analyses.

### Pollen quantification

To determine whether our diet treatments were effective, we quantified the amount of pollen consumed by lab-reared females receiving pollen, relative to reproductive females, by estimating the number of pollen grains in the hindgut. We followed previously described methods (Cane et al., 2016) to estimate pollen grains in 6 hindguts from each group. Individual hindguts were placed in 0.5 ml microcentrifuge tubes with 50 μl of 70% ethanol and torn apart with forceps. After guts were shredded, the mix was vortexed for 5 sec, using Vortex Genie 2 on highest setting of 10. The shredded gut was then removed using forceps, dabbing tissue on sides of the tube to remove excess ethanol and pollen. Each sample was vortexed on the highest setting for 10 seconds immediately prior to loading 10 μl of the solution into one chamber of a hemocytometer for pollen counting. Pollen grains were counted across the entire chamber under a compound microscope at 20X magnification. Three different 10 μl aliquots were counted for each sample, using the entire chamber each time. For each sample, the average of these three counts was divided by 0.0009 ml, the volume of each hemocytometer chamber, and then multiplied by the volume of ethanol used per sample (0.05 ml).

### Statistical analyses

All statistical analyses were performed in R version 3.2.5 (R Core Team, 2016). Visual inspection of a qq-plot (R package “car”, (Fox and Weisberg, 2011)) and an Anderson-Darling normality test (R package “nortest”, (Gross and Ligges, 2015)) revealed significant departures from normality in the distribution of maximum terminal oocyte and Dufour's gland lengths for Experiment 1, but not Experiment 2. We therefore applied a Box-Cox transformation to the data for Experiment 1 before running the final model (Venables and Ripley, 2002). For both Experiment 1 and 2, we modeled the maximum viable terminal oocyte and Dufour's gland lengths with separate linear mixed effects regressions that initially included intertegular width and treatment (coded as factors: diet for Experiment 1, JH*social for Experiment 2) as fixed effects, with bee bed of origin as a random effect (Bates et al., 2015). In each case, the variance and standard deviation for the intercept of bee bed was zero, so a linear model without random effects was used for subsequent analyses. Intertegular width was removed from the final models in the cases where it was non-significant (p > 0.05) – all except Dufour's gland length in Experiment 2. We used Tukey post-hoc tests to investigate significant differences between treatment groups (Hothorn et al., 2008).

For Experiment 2, we repeated the analyses after removing cases where the older, reproductive partner had a smaller intertegular width, maximum terminal oocyte, or Dufour's gland than the newly emerged cage-mate to determine whether relative size or reproductive development influenced the outcome of the social treatment.

We compared stages of oocyte maturation across treatment groups in Experiment 1 with ordinal logistic regression (R package “ordinal”, (Christensen, 2015)). However, this model was not appropriate for the data in Experiment 2, due to low representation of some values.

We compared the proportion of oocyte maturation stage among JH treatment groups in Experiment 2 using a chi-square test (R package “stats”, (R Core Team, 2016)). We also used a chi-square test to compare the proportion of samples with and without resorbed oocytes across groups in Experiments 1 and 2.

Final estimates of pollen counts in the hindgut were compared between groups (reproductive, nesting females, sugar & pollen mix, sugar, pollen, & alfalfa flowers) with a linear model function (lm), after applying a Box-Cox transformation of the data (Venables and Ripley, 2002).

## Results

### Patterns of ovary maturation

We used the reproductive and newly emerged females to describe patterns of oocyte maturation in alkali bees. Newly emerged bees activate their oocytes rapidly, even without mating. The modal stage of maturation for newly emerged females was one viable stage I oocyte, with a viable stage I or II oocyte in the other ovary. However, three newly emerged females had viable stage IV or V oocytes in both ovaries. It is possible that these females had spent a day or more in their nests upon eclosion, and were therefore slightly older than the others. Newly emerged females did not show any evidence of resorbing oocytes.

The modal stage of maturation for reproductive females was one viable stage IV oocyte and one viable stage IV or V oocyte in the other ovary. The maximum viable terminal oocytes in both ovaries were vitellogenic (stage III, IV, or V) for all reproductive females, except one with a pre-vitellogenic (stage II) oocyte in one ovary. Most (70%) reproductive females had at least one resorbing oocyte, and 35% had a resorbing oocyte in both ovaries. Resorbing oocytes were all in either category IVr or Vr, with 58% in stage Vr. The maximum viable terminal oocytes in ovaries with a resorbing oocyte were in stage III or IV, indicating that oocyte maturation is sequential across ovarioles within an ovary.

### Experiment 1 – nutrition effects on reproductive physiology

Mortality was not significantly different among lab-reared females on different diet treatments (mortality: sugar – 40%, sugar & pollen – 32%, sugar, pollen, & flowers – 46%; X^2^ = 1.29, p = 0.52, n = 92). There were significant differences in both maximum viable terminal oocyte and Dufour's gland length among treatment groups (oocytes: F4,89 = 30.68, r^2^ = 0.58, p < 4.79×10^−16^, Table S1; Dufour's: F4,101 = 45.80, r^2^ = 0.64, p < 2.20×10^−16^, Table S2). Among these groups, lab-reared, 10 d old females had significantly longer viable maximum terminal oocytes and Dufour's glands than newly emerged females (Fig. 1). However, actively nesting reproductive females had significantly more developed reproductive anatomy than either newly emerged or lab-reared females (Fig. 1). We did not observe significant differences in maximum viable terminal oocyte or Dufour's gland length among females reared in the lab for 10 d on different diets (Fig. 1).

**Figure 1.**
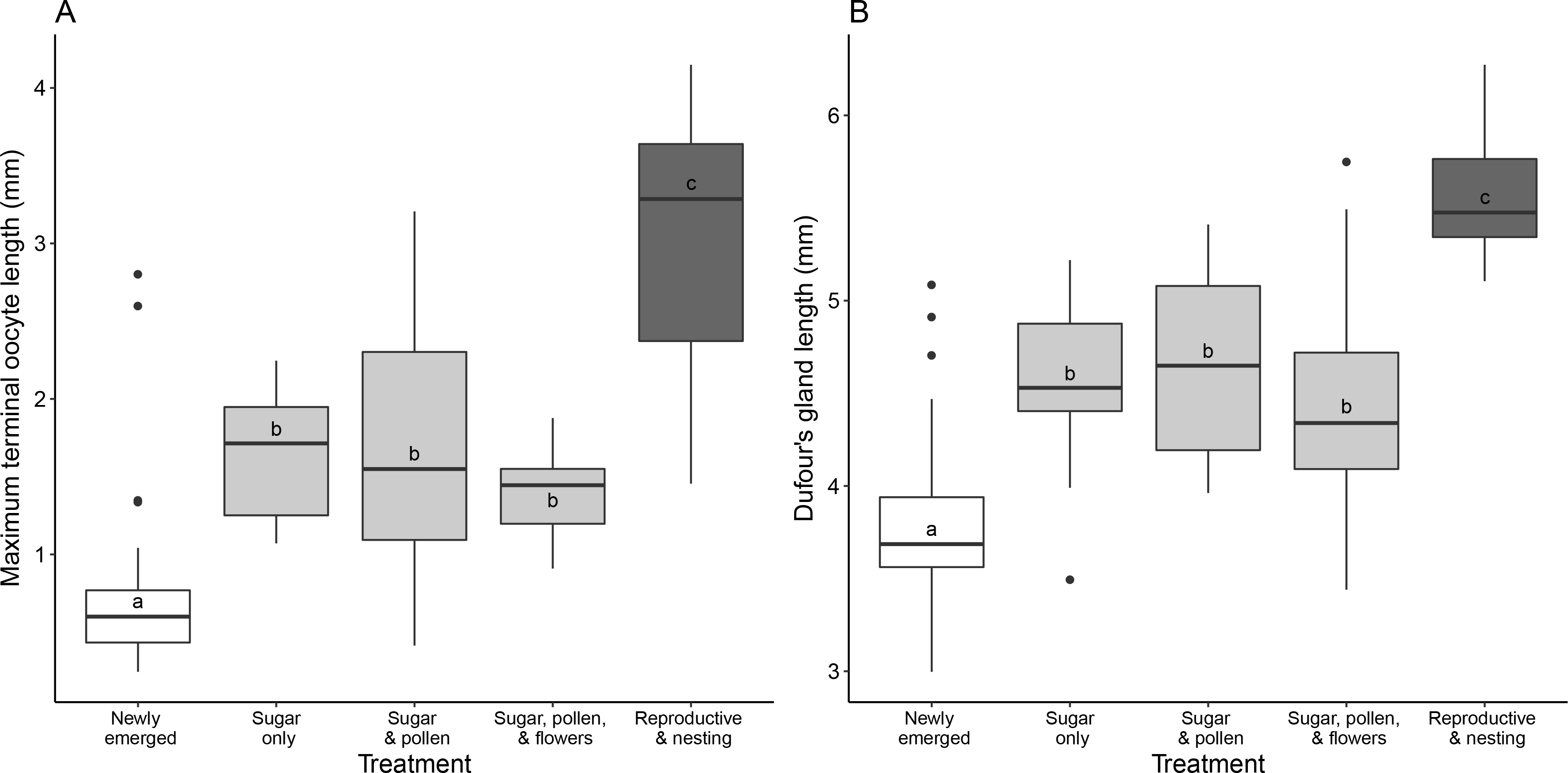
Effects of diet on reproductive maturation in alkali bees. (A) Maximum viable terminal oocyte length and (B) Dufour's gland length were significantly different between newly emerged, lab-reared, and nesting females (oocytes: F4,89 = 30.68, r^2^ = 0.58, p < 4.79×10^−16^; Dufour's: F4,101 = 45.80, r^2^ = 0.64, p < 2.20×10^−16^; n = individual bees, newly emerged – 36, sugar – 14, sugar & pollen – 22, sugar, pollen & alfalfa – 14, reproductive – 20). Diet did not have a significant effect on reproductive development when newly emerged females were reared in the lab for 10 d. Boxes represent the interquartile range, with the line as the median. Whiskers extend to 1.5 times the interquartile range. Circles are outliers. Letters indicate significant differences (p < 0.001 in Tukey post-hoc tests). White boxes = newly emerged females, Grey boxes = lab-reared 10 d old females; Dark grey boxes = Reproductive, nesting females; Full model results are available in the supplementary materials.

The maximum viable terminal oocytes of newly emerged females were at significantly lower stages of development than those of lab-reared 10 d old females or reproductive females, but there were no significant differences between females in the latter groups, all but one of whom had vitellogenic oocytes (stage III or higher) (ordinal logistic regression: Z = −4.61, p = 3.99×10^−6^; Fig. 2). Three lab-reared 10 d old females that received a sugar and pollen diet developed viable mature oocytes (stage V). Approximately half of the lab-reared females had at least one resorbing oocyte (sugar only – 50%, sugar & pollen – 50%, sugar, pollen, & flowers – 57%), and all except one (stage Vr) were in stage IVr. The proportion of females with resorbing oocytes among lab-reared females was not significantly different from that of reproductive females (X^2^ = 2.10, p = 0.55, n = 70 bees). However, most (58%) of the resorbing oocytes in reproductive females were in a more advanced stage (Vr).

**Figure 2.**
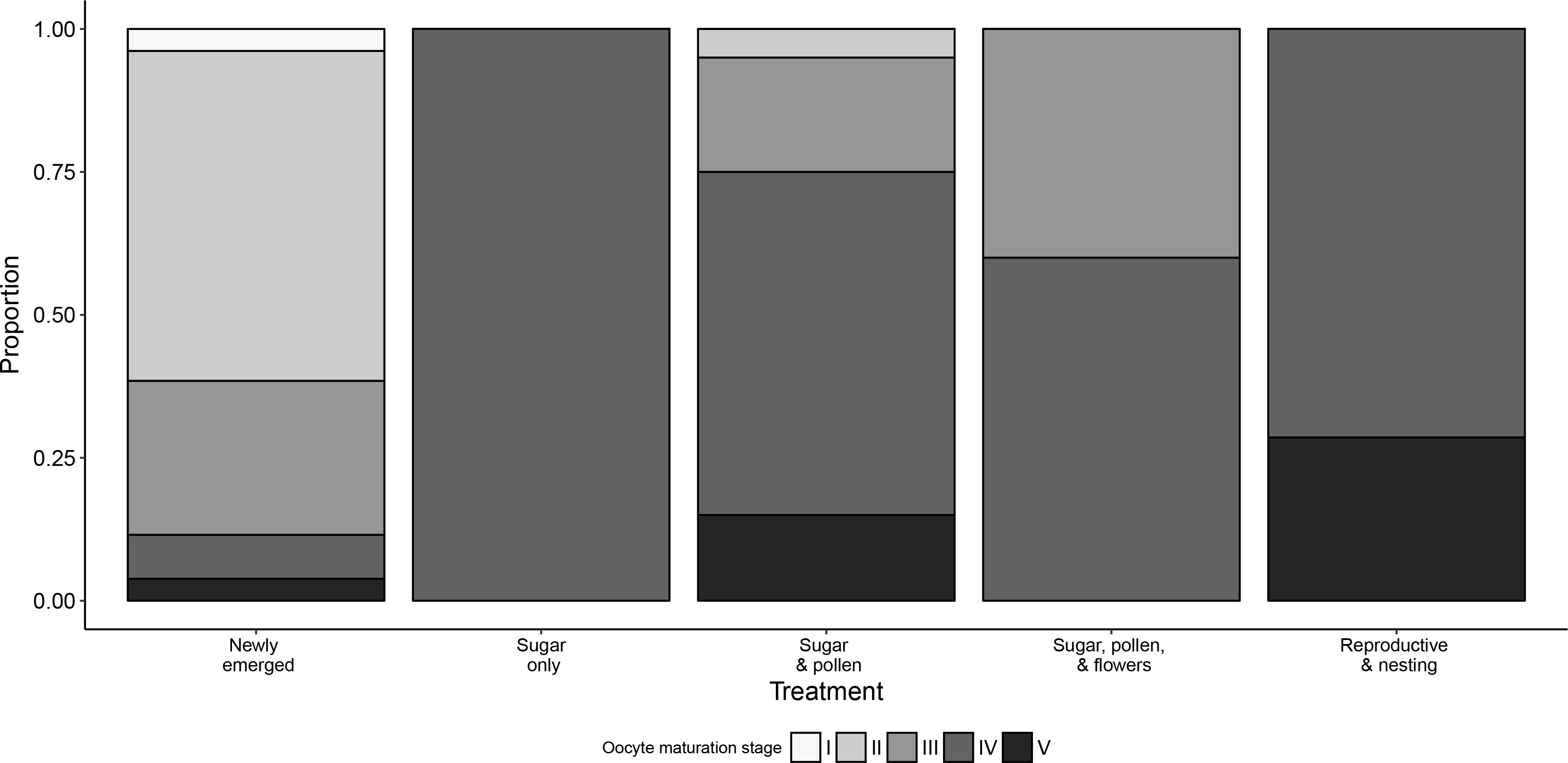
Effects of diet on oocyte maturation in alkali bees. Newly emerged females had oocytes at significantly earlier stages of maturation than females in the other treatment groups (ordinal logistic regression: Z = −4.61, p = 3.99×10^−6^, n = individual bees, newly emerged – 26, sugar – 11, sugar & pollen – 20, sugar, pollen & alfalfa – 10, reproductive – 7). Shading within bars indicates the proportion of maximum viable terminal oocytes in each stage of maturation, with stage I and II as pre-vitellogenic, III and IV as vitellogenic, and V as mature.

The estimated number of pollen grains detected in the hindguts was not significantly different among reproductive females and the two groups of lab-reared females that received pollen in their diet (F2,15 = 3.32, r^2^ = 0.31, p = 0.06, Fig. 3).

**Figure 3.**
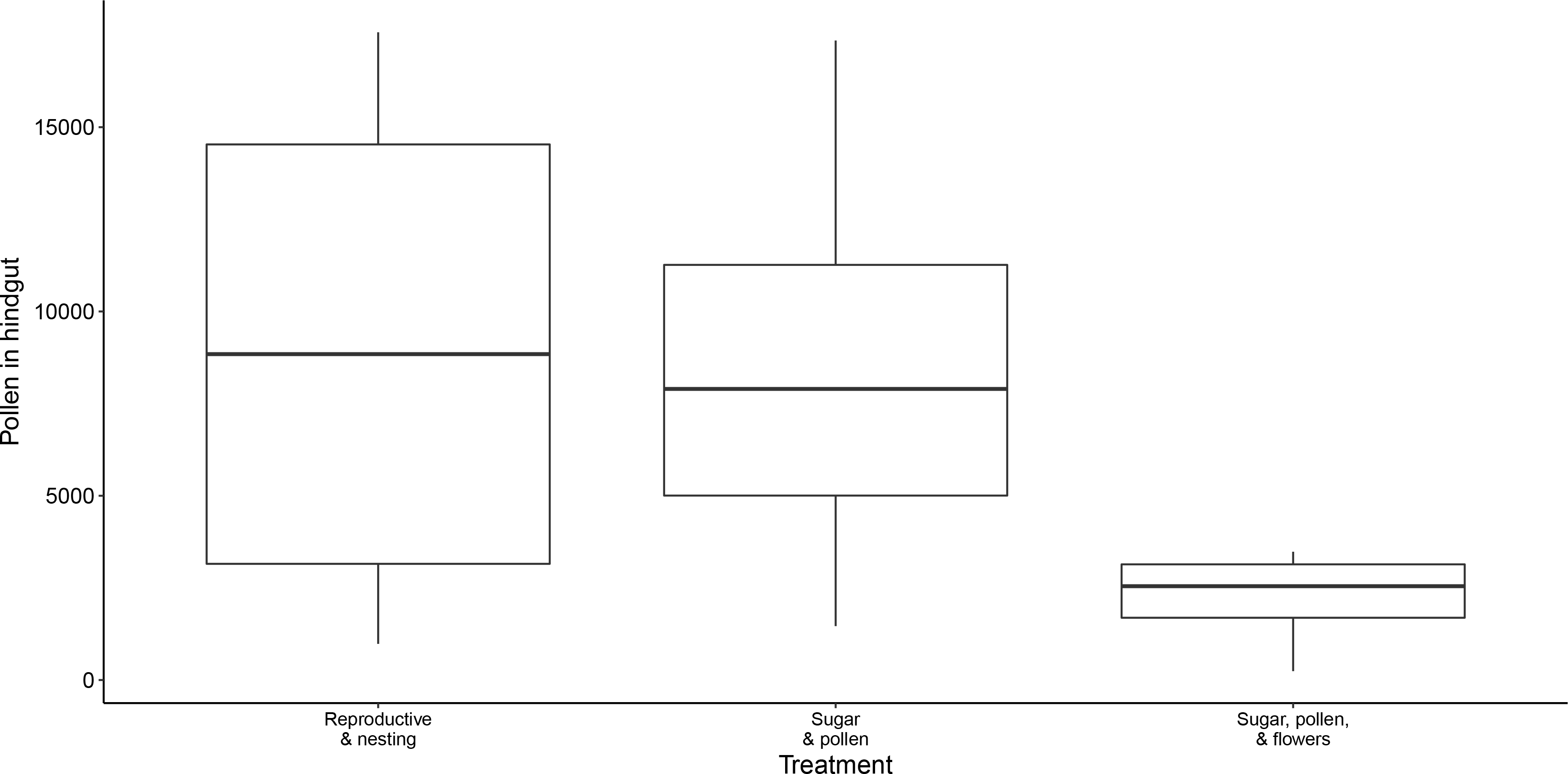
Estimates of pollen grains in the hindguts of female bees. There were no significant differences in hindgut pollen content between reproductive females and lab-reared females in posthoc tests (F2,15 = 3.32, r^2^ = 0.31, p = 0.06; n = 6 bees per group). Boxes represent the interquartile range, with the line as the median. Whiskers extend to 1.5 times the interquartile range.

### Experiment 2 – social and endocrine effects on reproductive physiology

Mortality was not significantly different among treatment groups in Experiment 2 (mortality: sham/solitary – 19%, sham/social – 19%, DMF/solitary – 20%, DMF/social – 47%, JH/solitary – 35%, JH/social – 44%; X^2^ = 7.04, p = 0.22, n = 101). There were significant differences in maximum viable terminal oocyte and Dufour's gland length among treatment groups (oocytes: F5,41 = 6.68, r^2^ = 0.45, p = 1.23×10^−4^, Table S3; Dufour's: F6,57 = 8.77, r^2^ = 0.48, p =8.97×10^−7^, Table S4). Females treated with JH had significantly longer viable terminal oocytes and Dufour's glands than females in control groups, but variation in the social environment did not have a significant effect on these measures of reproductive physiology (Fig. 4).

**Figure 4.**
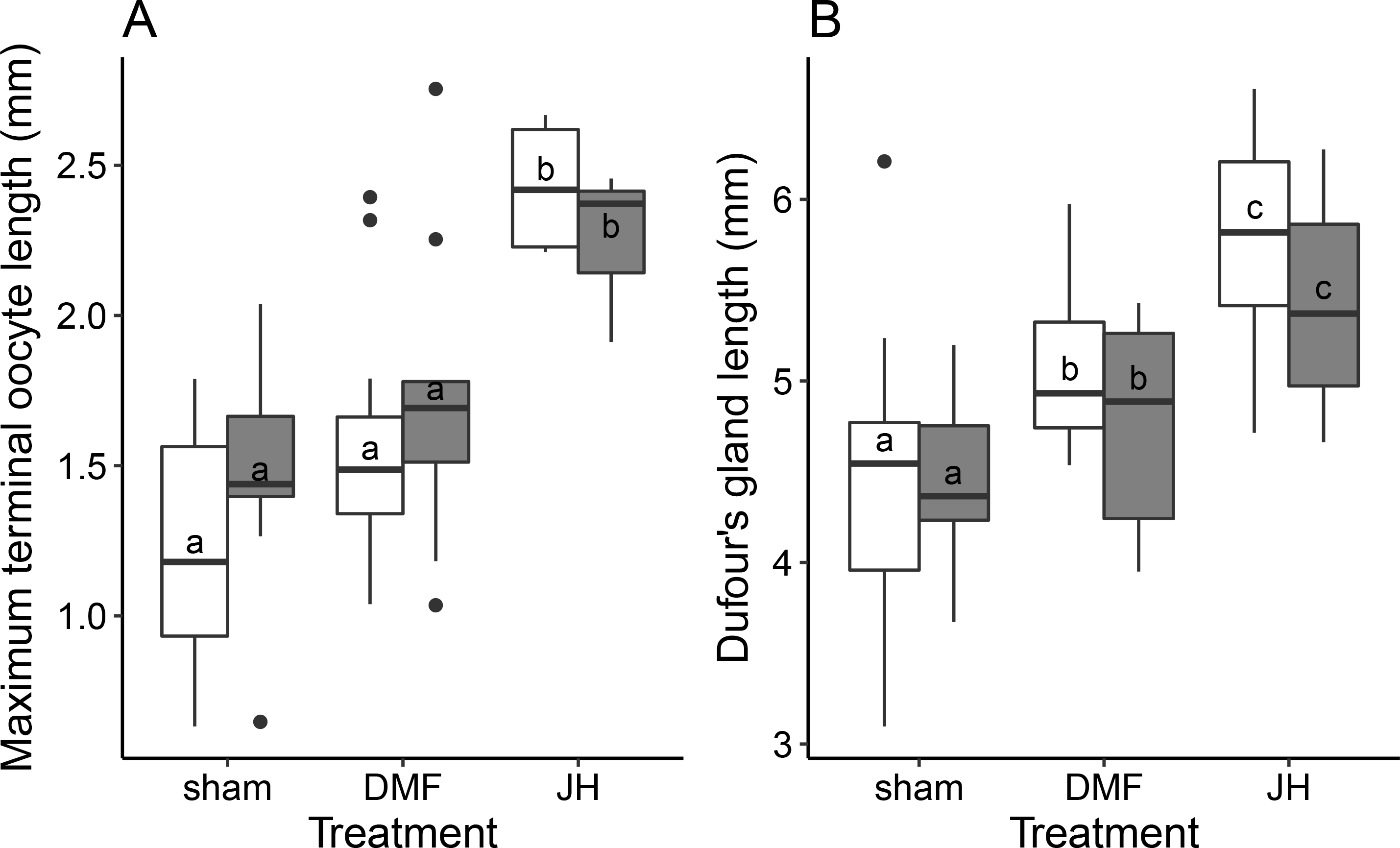
Effects of endocrine and social treatments on reproductive development in alkali bees. (A) Maximum terminal oocyte length and (B) Dufour's gland length were significantly different between lab-reared females treated with JH versus controls, but variation in the social environment did not significantly affect reproductive development (oocytes: F5,41 = 6.68, r^2^ = 0.45, p = 1.23×10^−4^; Dufour's: F6,57 = 8.77, r^2^ = 0.48, p =8.97×10^−7^); n = individual bees, sham/solitary – 12, sham/social – 12, DMF/solitary – 12; DMF/social – 10; JH/solitary – 10; JH/social – 8). Boxes represent the interquartile range, with the line as the median. Whiskers extend to 1.5 times the interquartile range. Circles are outliers. Letters indicate significant differences (p < 0.05 in Tukey post-hoc tests). Grey bars = social treatment, white bars = solitary treatment; Full model results are available in the supplementary materials.

Reproductive females and newly emerged females paired in the social treatments were similar in size to one another (mean ratio of intertegular width = 1.03 ± 0.11 s.d.). On average, the reproductive females had longer viable oocytes and Dufour's glands than their newly emerged cage-mates (mean ratio of maximum terminal oocyte length = 1.72 ± 0.62, mean ratio of Dufour's glands = 1.13 ± 0.17). However, there were 12 cases for which the reproductive female was smaller and/or had smaller ovaries or Dufour's glands than their newly emerged female cage-mates. Elimination of these 12 cases from the dataset did not change the results (Table S5-S6).

There were significant differences in stage of oocyte maturation among JH treatment groups (χ^2^ = 23.20, p = 7.31×10^−4^, n = 40 bees). Only females treated with JH had viable mature oocytes (stage V), and sham treated bees were the only group with pre-vitellogenic (stage II) oocytes (Fig. 5). Most (78%) of the JH-treated females had at least one resorbing oocyte, while 55-58% of the females in the control treatments had a resorbing oocytes. However, this difference was not statistically significant (X^2^ = 2.56, p = 0.28, n = 64). Resorbed oocytes were in stage IVr and Vr in each group.

**Figure 5.**
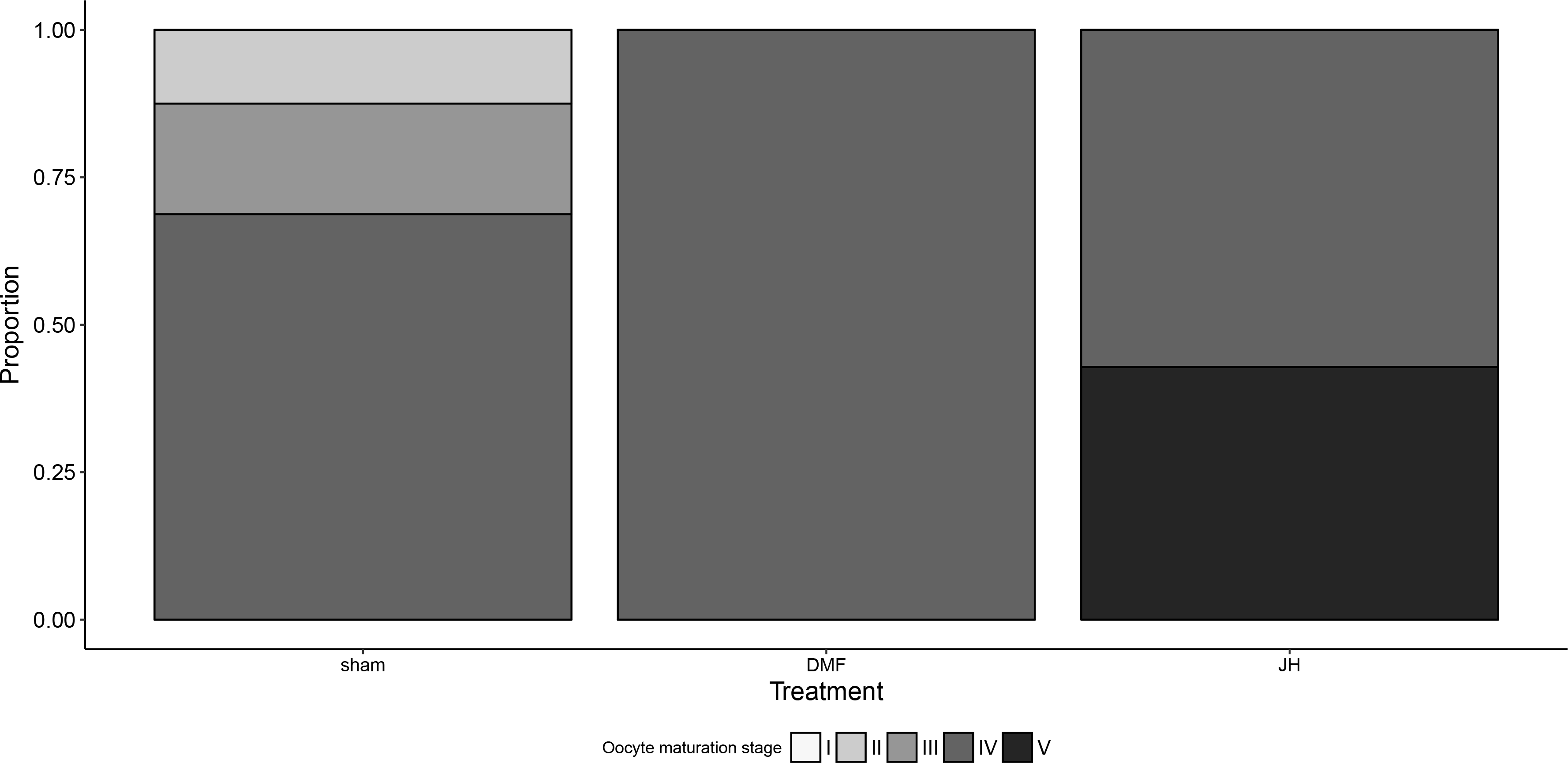
Effects of JH on oocyte maturation in alkali bees. There were significant differences in stage of oocyte maturation among JH treatment groups (X^2^ = 23.20, p = 7.31×10^−4^, n = individual bees, sham – 16, DMF – 17, JH - 7). Shading within bars indicates the proportion of maximum viable terminal oocytes in each stage of maturation, with stage I and II as pre-vitellogenic, III and IV as vitellogenic, and V as mature.

## Discussion

Variation in reproductive physiology is a hallmark of the social insect societies, in which just one or a few individuals out of thousands within a colony are reproductively active, despite shared genetic influences. Investigating the factors that contribute to reproductive variation in solitary relatives of social insects can provide clues as to how reproductive castes evolve (Kapheim, 2017). Our results demonstrate that solitary alkali bees do not require dietary protein during the initial stages of reproductive maturation, but that JH enhances this process. We also find that, unlike for social bees, interactions between conspecifics do not influence reproductive physiology. This provides important information about the physiological foundation from which reproductive castes emerged.

Access to dietary protein did not limit reproductive activation among newly emerged alkali bees, but it was rarely sufficient for reproductive maturation. Most of the lab-reared females in our study did not develop mature (stage V or Vr) oocytes or Dufour's glands during the 10 d study period, despite having similar amounts of pollen in their hindguts as actively nesting, reproductive females, indicating they had consumed ecologically relevant amounts of pollen during the experiment. Alkali bees commonly begin laying eggs within a few days of eclosion, and some of the newly emerged bees had mature oocytes, indicating that our study period provided ample time for reproductive maturation (Bohart and Cross, 1955). It is possible that the completion of reproductive maturation is hastened by ecological cues, such as nesting substrate, or access to mates. Seminal fluid is known to trigger oogenesis in several insect species (Avila et al., 2011), and mating limitation is known to influence reproductive activity in other halictid bees (Yanega, 1989; Yanega, 1992). However, lab-reared females of another halictid bee, *Megalopta genalis*, reached reproductive maturity when reared, unmated, in the lab for 10 days (Kapheim et al., 2012). If mating is a reproductive limitation in alkali bees, it can apparently be overridden by JH treatments, as JH treated females in our study were more likely to reach reproductive maturity, even in the absence of mating or ecological cues. Regardless of the role of dietary protein in the initiation of reproductive activation, the fact that alkali bees and other closely related halictid bees consume pollen on a daily basis suggests that protein is likely necessary for sustained reproductive activity throughout the breeding season (Cane et al., 2016; Wuellner, 1999).

Our results are in contrast to results of similar studies of solitary bees in the family Megachilidae, *Osmia californica* and *Megachile rotundata*, which showed access to dietary protein is essential for reproductive maturation among newly emerged females (Cane, 2016; Richards, 1994). Unlike alkali bees, *Osmia* overwinter as adults, begin oogenesis prior to eclosion, and thus eclose with depleted protein stores (Wasielewski et al., 2011). Protein stores have not been measured in newly emerged alkali bees, but remaining reproductively quiescent until eclosion is likely to be less energetically expensive and may be associated with increased availability of nutrient stores for post-eclosion maturation (Hahn and Denlinger, 2007). Alkali bees may therefore be better poised to initiate oogenesis without a dietary protein source. Conversely, pollen is necessary to stimulate vitellogenesis in *M. rotundata*, which also remains reproductively quiescent until ecolosion. The apparent difference in nutritional requirements for oogenesis among megachilid and halictid bees indicates that the physiological basis for reproductive activity is highly variable among solitary bees. This suggests that assumptions about reproductive physiology among the ancestors of social bees should be made with caution. Additional research on solitary bees from additional families in which eusociality has evolved (e.g., Apidae) are necessary to fully understand variation in nutritional requirements for reproduction.

Our results suggest that JH is a limiting factor in reproductive maturation among alkali bees. Females treated with JH were significantly more likely to have developed viable mature (stage V) oocytes and Dufour's glands, though other groups had mature resorbing (stage Vr) oocytes. This is consistent with earlier results (Kapheim and Johnson, 2017), and provides evidence of a conserved gonadotropic role for JH in alkali bees. In most insects, including bumble bees, JH stimulates the synthesis of vitellogenin, an egg-yolk precursor protein necessary for oocyte maturation (Amsalem et al., 2015; Badisco et al., 2013). This suggests the gonadotropic response to JH may depend on a dietary source of protein. All of the females receiving JH treatments in our study also received dietary protein from pollen, and thus had the nutritional resources necessary to complete vitellogenesis. Future studies are needed to determine how nutrition and JH pathways interact in alkali bee oogenesis.

The path by which JH stimulates Dufour's gland maturation is less clear, as Dufour's gland evolved in the ancestor of Hymenoptera, and secretes chemicals with a wide range of functions within this group (Mitra, 2013). Dufour's gland is likely derived from the colleterial accessory gland in other insects (Mitra, 2013), and reproductive maturity of this gland is induced by JH in cockroaches (*Byrsotria fumigate, Periplaneta americana*) (Bell and Barth, 1970; Willis and Brunet, 1966). Moreover, JH influences the chemical composition of Dufour's gland secretions in bumble bee (*B. terrestris*) workers (Shpigler et al., 2014). This, along with our results, suggests that endocrine regulation of Dufour's gland is deeply conserved among insects. Additional research is needed to determine the molecular mechanisms by which JH affects Dufour's gland function.

Unlike for social bees and some solitary bees, variation in the social environment does not influence reproductive physiology among solitary alkali bees. Most of the 20,000 species of bees are solitary, mass-provisioning, and annual (Michener, 2007). This means that after mating, females build a nest, provision each brood cell with pollen and nectar, lay an egg on top of those provisions, seal the cell, and die before any of her offspring complete development. For many of these species, a single mating event is the only interaction they have with conspecifics in their entire lives. Conversely, highly eusocial insects have evolved sophisticated forms of social communication. Within colonies of honey bees, bumble bees, and sweat bees, behavior and reproductive physiology is dynamically regulated by the behavior and/or pheromones of nestmates (Alaux et al., 2009; Amsalem and Hefetz, 2010; Arneson and Wcislo, 2003; Grozinger et al., 2003; Huang et al., 1998; Kapheim et al., 2016; Le Conte et al., 2001; Li-Byarlay et al., 2014; Padilla et al., 2016; Smith et al., 2009). Research with a facultatively eusocial halictid bee, *M. genalis*, suggests that aggression from older, reproductive females can limit reproductive development via JH-suppression in newly emerged females (Kapheim et al., 2016; Smith et al., 2009; Smith et al., 2013). However, the degree to which solitary bees respond to social cues varies among species. For example, experimental co-housing of otherwise solitary *Ceratina* bees (Family Apidae) results in division of labor and reproductive suppression in species that occasionally share nests in nature (Sakagami and Maeta, 1977; Sakagami and Maeta, 1984; Sakagami and Maeta, 1989; Sakagami and Maeta, 1995), but apparently not in species that never share nests, like alkali bees. Although we did not directly measure behavioral interactions among pairs as part of our study, we routinely observed aggressive exchanges among pairs of females in cages. Our results thus suggest that sensitivity to cues from the social environment observed in social halictid bees are not conserved in their solitary relatives. Alkali bees nest in extremely dense aggregations, with up to 713 nests per square meter in study area (Cane, 2008). At high density, these ground-nesting females are likely to encounter each other regularly as they dig tunnels and build cells, and there is thus likely to be strong selection against physiological sensitivity to social interactions in these populations. Similar studies with additional solitary bees are necessary to identify the circumstances under which sensitivity to the social environment influences reproductive physiology.

## Conclusions

This study is the first experimental investigation of dietary, endocrine, and social effects on reproductive maturation in a solitary bee closely related to lineages in which sociality evolved, but which is highly divergent (>100 my) from the more commonly studied eusocial honey bees and bumble bees (Cardinal and Danforth, 2013). Our results reveal that the factors contributing to the initiation of reproductive activation and completion of reproductive maturity may be different. Specifically, dietary protein was not essential for the initiation of reproductive activation, but was rarely sufficient for reproductive maturation. JH, however, may be a limiting factor in maturation of both oocytes and Dufour's gland. This provides important insight into how sensitivity to these cues evolved with the origin of reproductive castes in social insects. For example, the effects of JH on ovary and Dufour's gland maturation are apparently conserved between solitary alkali bees and bumble bee workers. However, these JH effects are responsive to social status in bumble bees, but not alkali bees (Amsalem et al., 2014; Shpigler et al., 2014). This suggests that different components of the endocrine networks influencing reproductive physiology were independently modified during social evolution. Also, nutrition and cues from the social environment are some of the most important factors in reproductive suppression of workers among social bees (Amsalem et al., 2013; Kapheim et al., 2016; Lawson et al., 2016; Padilla et al., 2016), but these factors did not have a significant influence on variation in reproductive activation in solitary alkali bees. This suggests that changes in how nutrient-sensing and environment-sensing pathways regulate reproductive physiology were especially important in the evolutionary origins of reproductive castes. Further comparisons of the molecular networks underlying the physiological response to nutritional, endocrine, and social cues across species are likely to provide key insight into how reproductive division of labor evolves.

## Acknowledgements

Tripodi for help with imaging spermathecae, E. Klinger for help with pollen counting, and J. Cane for helpful discussion. Thanks to A. Tripodi, D. Cox-Foster, and two anonymous reviewers for helpful comments on a previous version of this manuscript. We are grateful to John Dodd and Forage Genetics International for providing lab space and logistical support in Touchet, WA. We thank Mike Ingham, Mark Wagoner, and Mike Buckley for access to their bee beds and bees. M. Jolley and F. Dowsett provided valuable assistance in the field.

## Author contributions

KMK designed the study. KMK and MMJ conducted the experiments and analyses. KMK drafted the initial manuscript. KMK and MMJ revised the manuscript and approved the final version.

## References

Alaux, C., Le Conte, Y., Adams, H. A., Rodriguez-Zas, S., Grozinger, C. M., Sinha, S. and Robinson, G. E. (2009). Regulation of brain gene expression in honey bees by brood pheromone. Genes, Brain and Behavior 8, 309–319.

Amsalem, E., Grozinger, C. M., Padilla, M. and Hefetz, A. (2015). Chapter Two – The physiological and genomic bases of bumble bee social behaviour. In Advances in Insect Physiology, vol. Volume 48 eds. Z. Amro and F. K. Clement), pp. 37–93: Academic Press.

Amsalem, E. and Hefetz, A. (2010). The appeasement effect of sterility signaling in dominance contests among *Bombus terrestris* workers. Behavioral Ecology and Sociobiology 64, 1685–1694.

Amsalem, E., Shamia, D. and Hefetz, A. (2013). Aggression or ovarian development as determinants of reproductive dominance in *Bombus terrestris*: interpretation using a simulation model. Insectes Sociaux 60, 213–222.

Amsalem, E., Teal, P., Grozinger, C. M. and Hefetz, A. (2014). Precocene-I inhibits juvenile hormone biosynthesis, ovarian activation, aggression and alters sterility signal production in bumble bee (*Bombus terrestris*) workers. J Exp Biol 217, 3178–85.

Arneson, L. and Wcislo, W. T. (2003). Dominant-subordinate relationships in a facultatively social, nocturnal bee, *Megalopta genalis* (Hymenoptera: Halictidae). Journal of the Kansas Entomological Society 76, 183–193.

Avila, F. W., Sirot, L. K., LaFlamme, B. A., Rubinstein, C. D. and Wolfner, M. F. (2011). Insect seminal fluid proteins: identification and function. Annu Rev Entomol 56, 21–40.

Badisco, L., Van Wielendaele, P. and Vanden Broeck, J. (2013). Eat to reproduce: a key role for the insulin signaling pathway in adult insects. Front Physiol 4, 202.

Bates, D., Maechler, M., Bolker, B. and Walker, S. (2015). Fitting linear mixed-effects models using lme4. Journal of Statistical Software 67, 1–48.

Batra, S. W. T. (1970). Behavior of alkali bee, *Nomia-melanderi*, within nest (Hymenoptera-Halictidae). Annals of the Entomological Society of America 63, 400–406.

Batra, S. W. T. and Bohart, G. E. (1969). Alkali bees: response of adults to pathogenic fungi in brood cells. Science 165, 607.

Bell, W. J. and Barth, R. H. (1970). Quantitative effects of juvenile hormone on reproduction in the cockroach *Byrsotria fumigata*. Journal of Insect Physiology 16, 2303–2313.

Bohart, G. E. and Cross, E. A. (1955). Time relationships in the nest construction and life cycle of the alkali bee. Ann. Ent. Soc. Amer. 48, 403–406.

Brady, S. G., Sipes, S., Pearson, A. and Danforth, B. N. (2006). Recent and simultaneous origins of eusociality in halictid bees. Proceedings of Royal Society London B - Biological Sciences 273, 1643–9.

Cane, J. H. (1981). Dufour's gland secretion in the cell linings of bees (Hymenoptera: Apoidea). Journal of Chemical Ecology 7, 403–10.

Cane, J. H. (2008). A native ground-nesting bee (*Nomia melanderi*) sustainably managed to pollinate alfalfa across an intensively agricultural landscape. Apidologie 39, 315–323.

Cane, J. H. (2016). Adult pollen diet essential for egg maturation by a solitary *Osmia* bee. Journal of Insect Physiology 95, 105–109.

Cane, J. H., Dobson, H. E. M. and Boyer, B. (2016). Timing and size of daily pollen meals eaten by adult females of a solitary bee (*Nomia melanderi*) (Apiformes: Halictidae). Apidologie 48, 17–30.

Cardinal, S. and Danforth, B. N. (2013). Bees diversified in the age of eudicots. Proc Biol Sci 280, 20122686.

Christensen, R. H. B. (2015). ordinal - Regression Models for Ordinal Data.

Danforth, B. N., Eardley, C., Packer, L., Walker, K., Pauly, A. and Randrianambinintsoa, F. J. (2008). Phylogeny of Halictidae with an emphasis on endemic African Halictinae. Apidologie 39, 86–101.

Duchateau, M. J. and Velthuis, H. H. W. (1989). Ovarian development and egg laying in workers of Bombus terrestris. Entomologia Experimentalis et Applicata 51, 199–213.

Fox, J. and Weisberg, S. (2011). An R companion to applied regression. Thousand Oaks, CA: Sage.

Gibbs, J., Brady, S. G., Kanda, K. and Danforth, B. N. (2012). Phylogeny of halictine bees supports a shared origin of eusociality for *Halictus* and *Lasioglossum* (Apoidea: Anthophila: Halictidae). Molecular Phylogenetics and Evolution 65, 926–939.

Giray, T., Giovanetti, M. and West-Eberhard, M. J. (2005). Juvenile hormone, reproduction, and worker behavior in the neotropical social wasp *Polistes canadensis*. Proceedings of the National Academy of Sciences of the United States of America 102, 3330–3335.

Gross, J. and Ligges, U. (2015). nortest: Tests for Normality. In R package version 1.0-4.

Grozinger, C. M., Sharabash, N. M., Whitfield, C. W. and Robinson, G. E. (2003). Pheromone-mediated gene expression in the honey bee brain. Proceedings of the National Academy of Sciences of the United States of America 100, 14519–14525.

Hahn, D. A. and Denlinger, D. L. (2007). Meeting the energetic demands of insect diapause: nutrient storage and utilization. Journal of Insect Physiology 53, 760–773.

Hothorn, T., Bretz, F. and Westfall, P. (2008). Simultaneous inference in general parametric models. Biometrical J 50, 346–363.

Huang, Z. Y., Plettner, E. and Robinson, G. E. (1998). Effects of social environment and worker mandibular glands on endocrine-mediated behavioral development in honey bees. Journal of Comparative Physiology A: Neuroethology, Sensory, Neural, and Behavioral Physiology 183, 143–152.

Kapheim, K. M. (2017). Nutritional, endocrine, and social influences on reproductive physiology at the origins of social behavior. Current Opinion in Insect Science 22, 62–70.

Kapheim, K. M., Chan, T. Y., Smith, A. R., Wcislo, W. T. and Nonacs, P. (2016). Ontogeny of division of labor in a facultatively eusocial sweat bee *Megalopta genalis*. Insectes Sociaux 63, 185–191.

Kapheim, K. M. and Johnson, M. M. (2017). Support for the reproductive ground plan hypothesis in a solitary bee: links between sucrose response and reproductive status. Proc Biol Sci 284, 20162406.

Kapheim, K. M., Smith, A. R., Ihle, K. E., Amdam, G. V., Nonacs, P. and Wcislo, W. T. (2012). Physiological variation as a mechanism for developmental caste-biasing in a facultatively eusocial sweat bee. Proceedings of the Royal Society B: Biological Sciences 279, 1437–1446.

Lawson, S. P., Ciaccio, K. N. and Rehan, S. M. (2016). Maternal manipulation of pollen provisions affects worker production in a small carpenter bee. Behavioral Ecology and Sociobiology 70, 1891–1900.

Le Conte, Y., Mohammedi, A. and Robinson, G. E. (2001). Primer effects of a brood pheromone on honeybee behavioural development. Proceedings of the Royal Society of London. Series B: Biological Sciences 268, 163–168.

Li-Byarlay, H., Rittschof, C. C., Massey, J. H., Pittendrigh, B. R. and Robinson, G. E. (2014). Socially responsive effects of brain oxidative metabolism on aggression. Proceedings of the National Academy of Sciences.

Michener, C. D. (1974). The social behavior of the bees. Cambridge, MA: Harvard University Press.

Michener, C. D. (2007). Bees of the world. Baltimore, MD: The Johns Hopkins University Press.

Mitra, A. (2013). Function of the Dufour's gland in solitary and social Hymenoptera. Journal of Hymenoptera Research 35, 33–58.

Oliveira, R. C., Vollet-Neto, A., Akemi Oi, C., van Zweden, J. S., Nascimento, F., Sullivan Brent, C. and Wenseleers, T. (2017). Hormonal pleiotropy helps maintain queen signal honesty in a highly eusocial wasp. Sci Rep 7, 1654.

Padilla, M., Amsalem, E., Altman, N., Hefetz, A. and Grozinger, C. M. (2016). Chemical communication is not sufficient to explain reproductive inhibition in the bumblebee *Bombus impatiens*. R Soc Open Sci 3, 160576.

R Core Team. (2016). R: A language and environment for statistical computing. Vienna, Austria: R Foundation for Statistical Computing.

Richards, K. W. (1994). Ovarian development in the alfalfa leafcutter bee, *Megachile rotunda*. Journal of Apicultural Research 33, 199–203.

Roulston, T. H. and Cane, J. H. (2000). Pollen nutritional content and digestibility for animals. Plant Systematics and Evolution 222, 187–209.

Sakagami, S. F. and Maeta, Y. (1977). Some presumably pre-social habits of Japanese *Ceratina* bees, with notes on various social types in Hymenoptera. Insectes Sociaux 24, 319–343.

Sakagami, S. F. and Maeta, Y. (1984). Multifemale nests and rudimentary castes in the normally solitary bee *Ceratina-japonica* (Hymenoptera, Xylocopinae). Journal of the Kansas Entomological Society 57, 639–656.

Sakagami, S. F. and Maeta, Y. (1989). Compatibility and incompatibility of solitary life with eusociality in two normally solitary bees *Ceratina japonica* and *Ceratina okinawana* (Hymenoptera, Apoidea), with notes on the incipient phase of eusociality. Jap J Entomol 57, 417–739.

Sakagami, S. F. and Maeta, Y. (1995). Task allocation in artificially induced colonies of a basically solitary bee *Ceratina* (*Ceratinidia*) *okinawana*, with a comparison of sociality between *Ceratina* and *Xylocopa* (Hymenoptera, Anthophoridae, Xylocopinae). Jap J Ecol 63, 115–150.

Schwarz, M. P., Richards, M. H. and Danforth, B. N. (2007). Changing paradigms in insect social evolution: insights from halictine and allodapine bees. Annual Review of Entomology 52, 127–50.

Shpigler, H., Amsalem, E., Huang, Z. Y., Cohen, M., Siegel, A. J., Hefetz, A. and Bloch, G. (2014). Gonadotropic and physiological functions of juvenile hormone in bumblebee (*Bombus terrestris*) workers. PLoS One 9, e100650.

Smith, A. R., Kapheim, K. M., O’Donnell, S. and Wcislo, W. T. (2009). Social competition but not subfertility leads to a division of labour in the facultatively social sweat bee *Megalopta genalis* (Hymenoptera: Halictidae). Animal Behaviour 78, 1043–1050.

Smith, A. R., Kapheim, K. M., Perez-Ortega, B., Brent, C. S. and Wcislo, W. T. (2013). Juvenile hormone levels reflect social opportunities in the facultatively eusocial sweat bee *Megalopta genalis* (Hymenoptera: Halictidae). Horm Behav 63, 1–4.

Venables, W. N. and Ripley, B. D. (2002). Modern applied statistics with S. New York: Springer.

Wasielewski, O., Giejdasz, K., Wojciechowicz, T. and Skrzypski, M. (2011). Ovary growth and protein levels in ovary and fat body during adult-wintering period in the red mason bee, *Osmia rufa*. Apidologie 42, 749–758.

Wcislo, W. T. and Engel, M. S. (1996). Social behavior and nest architecture of nomiine bees (Hymenoptera: Halictidae; Nomiinae). Journal of the Kansas Entomological Society 69, 158–167.

Willis, J. H. and Brunet, P. C. J. (1966). The hormonal control of colleterial gland secretion. Journal of Experimental Biology 44, 363–378.

Winston, M. L. (1987). The biology of the honey bee. Cambridge, MA: Harvard University Press.

Wuellner, C. T. (1999). Alternative reproductive strategies of a gregarious ground-nesting bee, *Dieunomia triangulifera* (Hymenoptera: Halictidae). Journal of Insect Behavior 12, 845–863.

Yanega, D. (1989). Caste determination and differential diapause within the first brood of *Halictus rubicundus* in New York (Hymenoptera, Halictidae). Behav Ecol Sociobiol 24, 97–107.

Yanega, D. (1992). Does mating determine caste in sweat bees - (Hymenoptera, Halictidae). Journal of the Kansas Entomological Society 65, 231–237.

